# Utilizing Quantum Biological Techniques on a Quantum Processing Unit for Improved Protein Binding Site Determination

**DOI:** 10.1101/2020.03.20.000950

**Authors:** Samarth Sandeep, Vaibhav Gupta, Torin Keenan

**Affiliations:** Iff Technologies LLC

## Abstract

Iff Technologies has constructed a tool named Polar+ that can predict protein-to-protein binding sites on a given receptor protein that operates faster and at a higher quality than the prominent industry standards for protein binding, including Autodock Vina and SwissDock. The ability to provide this advantage comes from a new approach to biophysics, dubbed many-body biological quantum systems, that are modeled using quantum processing units and quantum algorithms. This paper provides both experimental and theoretical evidence behind the validity of the quantum biology approach to protein modeling, an overview of the first experimental work completed by Polar+, and a review of the results obtained.

## I. INTRODUCTION

Succinctly modeling protein binding is essential for understanding the interactivity of different biological systems between one another, for designing impactfully new biological compounds, and for building biomaterials that combine both biological systems and non-biological systems, such as crystalline and amorphous systems. This has been made especially important recently by the advent of gene therapeutics’ promises of curing various diseases that has been further amplified through the development of CRISPR.

Precise modeling is still difficult because of the complexity associated with each amino acid. While there are many exacting models describing a protein’s interaction with other proteins or within certain solvents, it is still incredibly difficult to also model the effects of temperature, the effects of perturbations caused by other systems within the same area, and others[1].

If one were to assign function classes to this entire function, the function for binding becomes exponentially more difficult to solve as the number of amino acids increases, making it infeasible for computers that can only work in polynomial time to effectively solve for every possible combination.

There have been two classes of solutions to this problem. One class involves the use of machine learning and artificial intelligence in the space of biological modeling. This requires an exponential model to be approximated with a polynomial one, and would use large data sets of protein information and high performance clustering to recognize the similarities and differences between the analyzed protein and the test data. While this has seen huge success, especially through the demonstration greater precision in binding models, the data intensity of the models means that the machine learning is only precise as the data fed to it. While this is not a problem for other areas that machine learning has impacted, such as in speech recognition, where data sets are readily available for many types of speech patterns, protein modeling still depends on nanoscale imaging and assay techniques to characterize docking sites and linkages between amino acids, exacerbating the expense of machine learning for protein binding determination. Recently, other organizations have developed quantum algorithms to help increase the speed and precision of machine learning algorithms. This is part of the growing interest amongst machine learning researchers to experiment with quantum circuits with the goal of dynamically improving their classifiers by being able to evaluate a formula’s multiple parameters at a fraction of the time[2]. This has tremendous potential to increase both the speed and precision of protein binding as more combinations can be tried at the same time, but would still require data stores for each protein to be thorough.

However, all of these methods are based on the theory that amino acids in proteins interact with each other in a classical manner, and that the entire process is just sequential and highly parallelized in continuous operation of living systems[3]. But the speed, directional change, and magnetic fields of these interactions indicate a quantum effect that could occur between and within macroscopic biological systems.

Over the past decade, a large body of work has been completed to prove these quantum biology effects in a variety of different forms and at different scales, ranging from models of electron clouds in DNA to new theories around metabolic pathways. Thus, with this newfound interpretation of how biological systems actually work, the best way to model these systems must also take into account the quantum interactions within each system.

Polar+ is the first platform to take quantum biological modeling, apply it to a quantum computer, and treat the quantum computer as an analog for a protein. As such, it provides an advantage over all competitors, classical or quantum, in speed, precision, and cost.

This paper reviews how a quantum mechanical model of proteins simulated using a quantum computer can provide a distinct advantage over all other protein binding implementations. As such, the paper serves to answer two questions:

1. Can a protein be effectively modeled using super-conducting Josephson Junctions?
2. What is the improvement of using a quantum mechanical model for binding over other algorithms?

### A. Part 1: Using Josephson Junctions to Model Proteins

Using quantum mechanical models to describe biological systems is not novel; one of the foremost conferences regarding quantum biology in all history, The Versailles Conference on Theoretical Physics and Biology, was held in 1967, with some of the most impactful work in the field published around the same time[4]. However, much of this work was theory, and existed before the tools for proving quantum mechanical interactions, both mathematically and experimentally, were invented. The most well-known example of early quantum biology work is Lowdin’s work around DNA mutations[5], which states that the movement of protons across different amino acids in a protein to create mutations is a result of proton tunneling to reach an energy minimum or maximum, rather than a sequentially static process between two points.

But, as mathematical tools for describing quantum systems have become more widely used across disciplines, especially with Hartree-Fock chemical models[6], quantum biology research began to reemerge, with conferences and research groups dedicated to the study occurring world-wide, especially over the past decade. This was most apparent with Phillip Kurian’s[7] and Elisabeth Rieper’s work in endonuclease splitting[8], which characterized a Hamiltonian to describe the energy across a strand of DNA given a set of *N* nucleotides:

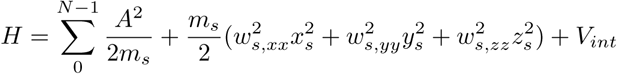

can be rewritten in terms of polar coordinates as:

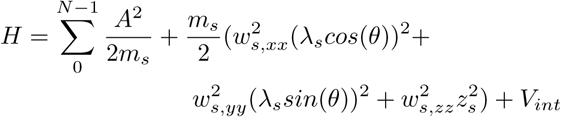

where λ refers to the distance between frequency events, which can be seen as the functions wavelength

While this model was developed for a helical structure in mind, the Hamiltonian is evaluated on a “rung-by-rung” basis, which means that it can be applied to other geometries as well, such as a protein that consists of more variations of the same nucleotides than exist within DNA, where A is equal to the distance from equilibrium and x, y, and z represent the difference in distance of each axis between the electron cloud

The live proof for this Hamiltonian has been observed by a few different sources. Most notably, an effort from Erling Thyrhaug [9] shows the correlation between a theoretical quantum system and biological interactions within a cellular system built off of this Hamiltonian further.

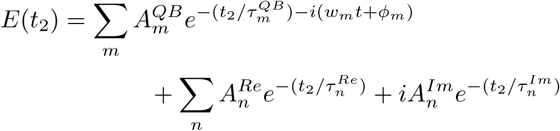

Where QB represents quantum beat, Re represents the real aspect of the wave, and Im represents the imaginary part of the wave. However, the Re and Im parts really represent a damping adjustment on the quantum beats that exist in the Fenna Matthews Olson (FMO) complex, a cohesive grouping of pigments and proteins found in bacteria, as the experimental model is based on the premise that the entire system is an underdamped circuit. Thus, taking what’s remaining of the FMO equation for energy between the different biological molecules, it can be rewritten in terms of sine and cosine as:

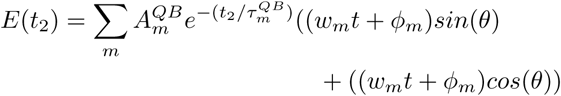

Further experimental evidence from the past decade strongly suggests that utilizing the Rigetti system for modeling of these quantum biological systems is appropriate. In November 2019, Armin Shayeghi and a team at the University of Vienna proved that gramicidin, an amino acid within some proteins, behaves in a quantum mechanical manner, with amino acids interacting with one another over magnetic fields with a positively-correlated Wigner function[10].

Additionally, there has been more interest in the quantum behavior of pigments, with a specific focus on cryptochrome[11], a protein that is found in avian and mammalian retinae that is believed to be a strong magnetoreceptor utilizing quantum transport that helps to guide avian direction.

Given this Hamiltonian interpretation, a chain of amino acids can be related to a quantum system. Moreover, it can directly relate to a macroscopic, many-bodied quantum system, such as a Josephson Junction, but more specifically a noisy Josephson Junction, who’s Hamiltonian is defined here:

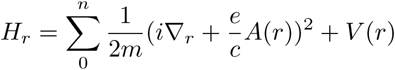

While these are equivalent and a Josephson Junction in theory could represent these biological systems, the actual setup of the Rigetti superconducting circuits needs to be evaluated to see if their system is a good fit for biomolecular modeling. Based on the work that Chad Rigetti[12] had completed leading up to the creation of the Rigetti Aspen 4Q used in this study, the superconducting circuit he had designed followed a circuit Hamiltonian of this form:

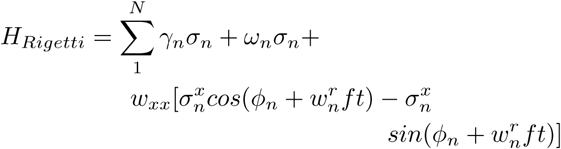

Thus, a multiple qubit system built using Rigetti’s architecture should be a perfect analog of the energy interactions that exist within a protein’s amino acids. The only issue is that of noise. All quantum models for biological systems highlight the noisiness of the systems that each of them operate in, regardless of coherence and decoherence; noiseless systems are unusual, but for practical purposes can be described with an updated Heisenberg limit provided that parameterization of priors is flat and no randomness is assumed, or at least if noise is eliminated using quantum error correction techniques[13], techniques which are not documented or used in this work. This is particularly highlighted in Filippo Caruso’s work on noise in quantum biological systems, and in particular the FMO complex [14]. The dephasing for this complex was found to be:

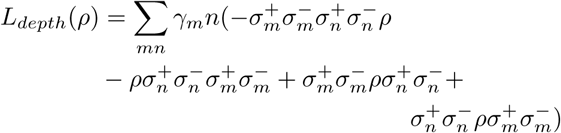

Where +/- represents operators that raise and lower the energy states of the given sites m and n and represents the current density between the two sites. According to Rigetti’s documentation [15], the noise on their qubits can be modeled by:

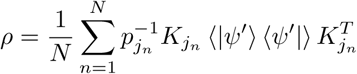

in which *p* is equal to probability, *K* is equal to a +/- operator depending on state, *ψ* represents the expectation value, and *N* represents the number of qubits. Therefore, since the 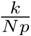 performs the same function as the sigma in the FMO equation, the *ψ* represents *γ* in the first equation and both are equivalent. Thus, the Rigetti Quantum Processing Unit should be able to serve as the perfect analog for a biological system’s modeling.

### B. Part 2: Improvements of a Quantum Mechanical Model for Modeling over Classical models

#### 1. The Best in Protein Binding Modeling

Protein binding is usually broken up into the categories of protein docking and protein-to-protein binding. Protein docking involves the connection of a protein to a ligand, which is a biomolecule that provides a signal when bound to a protein in the form of the distribution of some resource or some charge, while protein-to-protein binding usually involves complete protein chains binding to one another to produce a more complicated energy pathway with a cell. While protein-to-protein binding is important, understanding protein docking from a computational perspective is non-trivial, as it can be the core work performed by a multitude of lab tests when searching for new proteins or ligands for different use cases. It is for this reason that protein docking software is one of the most commonly used computational biology tools.

The most prominent of these tools is AutoDock [16]. Having existed for 30 years, Autodock has established itself as the industry standard for binding, and has made consistent updates to stay at the cutting edge. Most notably, the last major update was provided with the launch of the Vina platform in 2013, which provided significant improvements to the scoring function used within Autodock for calculating binding affinities by employing “force fields” and more algorithmic work beyond geometric modeling. Other algorithms, such as CBDock, use similar methods for modeling, with SwissDock using a grid based system and a rDock using a sphere based system to help identify potential binding sites [16].

While the performance that each algorithm can provide is definitely improved, there are little to no adjustments to changes of solvent, to changes due to temperature, to free energy perturbations, or to magnetic effects. The researcher is put in charge of determining these characteristics before or after running a binding test, which results in many researchers needing to run many trials and being unable to completely model protein interaction *in silico*.

Menten AI [17] and ProteinQure [18] both provide novel ways of using quantum computers in biological modeling work. While Menten’s work focuses more on sidechain additions to a protein and the creation and expression of peptides, ProteinQure’s work focuses on protein-ligand docking. ProteinQure’s solutions focus only on pharmacore points, which are pre-screened points-of-importance to pharmaceutical processes, rather than the entire protein’s binding site mapped for other modeling purposes. Their sampling methods are able to provide a speedup to the classical methods for modeling these points. However, they do not take into account the quantum nature of the biological macromolecules. Thus, the precision and universality of their binding points are still approximations of a much more complicated structure, rather than exact simulations of the entire proteomic systems.

#### 2. Polar+

The Polar+ system for protein binding modeling improves on current binding affinity algorithms by better defining the best and most polarized binding sites that exist within a protein. This allows for protein binding modeling beyond ligand binding, and can present a way to improve protein modeling universally. For the purposes of comparison, Polar+ will be reviewed for its ability to produce protein maps that can improve results in protein-ligand binding models.

Polar+ achieves an improvement in protein modeling by utilizing pre-processing of protein geometries from an atom-by-atom perspective, with binding site calculations through simulation of amino acid using qubits, and post-processing that effectively takes the final site calculations into account when presenting a final set of binding sites. In order to effectively compare Polar+ to the industry leaders, Polar+ will be run in tandem with Swiss-Dock, CBDock, RDock, and Autodock Vina over a given protein. These results will then be compared to results gained in the lab for actual protein binding observed. In order to effectively compare Polar+’s speeds to similar products, its speed will be compared to RDock and to classical protein binding site modeling using *k* –nearest neighbors clustering.

#### 3. Experimental Setup

As a first test of the advantage that a direct analog, the quantum computer, can provide over classical algorithms, the Iff team developed an algorithm for mapping the best potential binding sites within a protein.

The algorithm used to complete this reduction of sites was the Quantum Approximate Optimization Algorithm(QAOA). Given a graph of nodes *N*, the algorithm applies an Ising Hamiltonian onto an abstracted graph of *N* qubits and uses current density between these qubits to decide the cutting points of the graph [19].

In order to utilize this algorithm for a graph of atoms that make up a protein, pre-processing was completed to map sets of atoms with similar arrangements to the qubits within the 13 qubit Aspen 4Q-13E quantum computer provided by Rigetti, the only quantum computer with a geometry most adaptable to the different structures within the protein.

Once the mapping is completed, the splitting process can begin within the quantum computer. The outputs of this process are a set of bitstrings that correspond to different combinations of cuts, as described in Rigetti’s documentation. Using these cuts, one can find both redundant and energetically unfavorable binding sites, and eliminate them.

**FIG. 1.**
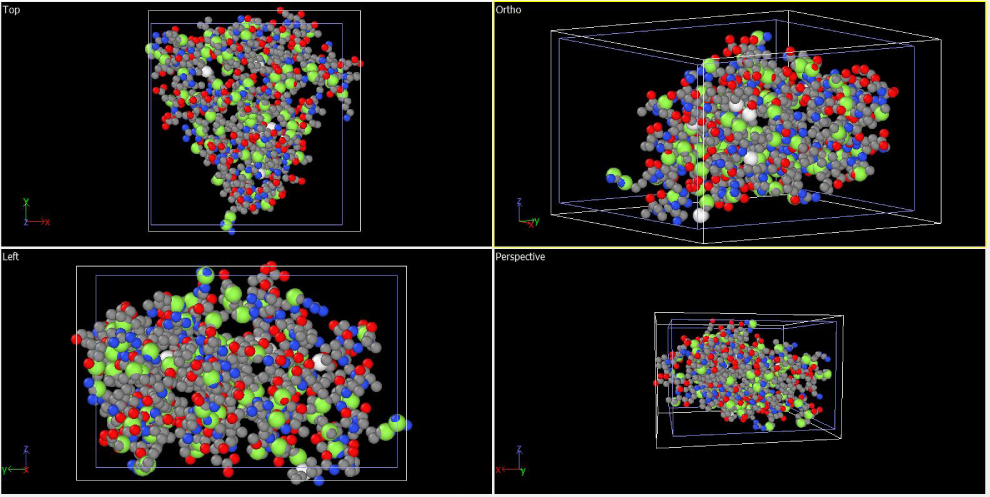
Binding sites within the protein 4Q2S. The Green are the Polar+ binding sites

**FIG. 2.**
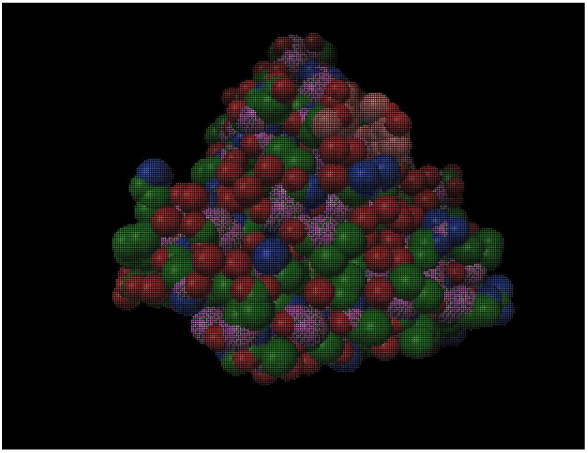
Binding sites within the protein 4Q2S interfacing with Resveratrol. The purple are the Polar+ binding sites. The salmon are the Resveratrol sites. The rest of the figure is the 4Q2S structure

For this experiment, the 4Q2S protein, one of the largest receptor proteins within the yeast cytoplasm was mapped, and it’s top binding sites were identified. To verify the validity of these binding sites, they were utilized to find potential binding conformations with the ligand, resveratrol.

#### 4. Experimental Results

After running this algorithm 4 times, the binding sites obtained were the same for each time the algorithm was run. The average run time for the algorithm on the QPU was 1 minute and 17 seconds.

While the average time taken to run this algorithm is much longer and slower than the times obtained for any of the docking softwares, a simple comparison between them does not take into account the level of complexity that QAOA can provide that docking softwares cannot. While docking software is focused mainly on geometric configurations and is not at all times for energy perturbations, QAOA can provide a probabilistic distribution of potential energy values that can be utilized for far more binding models beyond protein to protein binding. The only algorithms, then, that can effectively be a classical measure against QAOA in this use case would have to be a probabilistic energy function for each and every atom. GROMACS [20] is currently the leader in this area. However, the average run time on an 8-core machine for their algorithm is 5-6 hours. Thus, not only does Polar+ offer more precise information, it does so in a faster time.

### C. Complexity Class Comparisons

Most notably, the major difference that exists between modeling proteins and their binding sites on a quantum computer versus on any classical machine is the difference in complexity classes available to solve within. Looking at the most recently well-cited model for statistical modeling and classifications of proteins by Dr. Susanne Gerber at the University of Mainz[21], the highest degree of complexity achieved was a 3rd Degree polynomial time *O(n)* with bounds in *Z*. While this seems to be sufficient for a summation with 2 bounds, the complexity of completing this summation for an entire protein quickly becomes factorial, as each functional addition brings about another set of parameters to work through, as described here for just the kinetic energy of a protein, without taking into account the different material affects with each solvent.

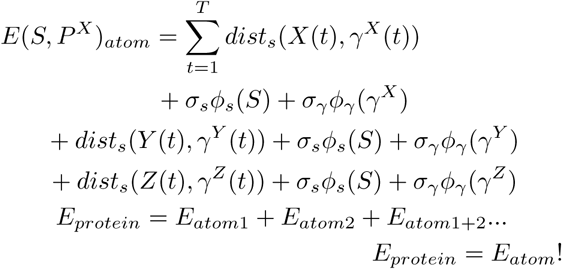

as every one of the internal terms is made factorial

Thus, with the Rigetti quantum computer really as an analog for this system, the calculations for these energy states do not need to be completed; rather, the quantum computer would complete energy transformations according to gate transformations.

The exact amino acids to which these energetics belong to would be reaffirmed in a post-selection process, which would only need to be done in polynomial time. Therefore, not only would the quantum calculation actually be completed in 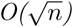 space due to an effective use of the hadamard gates, but the algorithmic work would be completed in *O(n)* time.

### D. Conclusion

This paper successfully demonstrates an advantage of a Rigetti-enabled algorithm over a classically demonstrated algorithm in the use case of protein binding. By identifying the most optimal binding sites within a protein, Polar+ running on the Rigetti Aspen 4Q-13E was able to provide more detail at a lower time than achieved classically.

However, the QAOA algorithm is really not the perfect algorithm for the modeling of quantum systems. In this use case, QAOA still treats the different protein binding sites as classical objects of set states, and uses a quantum algorithm between them to find the most optimal sites. But, this could be vastly improved if each binding site were instead initialized with a quantum wave function, rather than a set, classical state.

Moreover, the experimental setup could have greatly improved. Access to the QPU was obtained through the use of the Quantum Machine Image on top of a Amazon Web Services (AWS) EC2 Container. This level of abstraction away from the host computer could have resulted in a decrease in the speed obtained over a more direct setup with the QPU.

Therefore, while the results obtained through this experiment were encouraging and do provide an improvement to protein docking techniques, there still exist a multitude of unexplored experimental setups that could lead to a further advantage when truly utilizing the ability of the Rigetti machine to act as an analog to biological macromolecules.

## E. Acknowledgements

The Iff team is extremely grateful for the feedback and direction in protein modeling provided by Dr. Nitin at the Food Science department at the University of California, Davis; for further insight into the world of quantum biology provided by the *Quantum Spins in Biology* conference at the University of California, Los Angeles and its director, Dr. Clarice D. Aiello; for input into complexity class modeling provided by the *Quantum Wave in Computing Bootcamp* at the Simons Institute, University of California, Berkeley; and, last but not least, for continuous feedback and paper review from Slack user Moogoo.

